# CAGEd-oPOSSUM: motif enrichment analysis from CAGE-derived TSSs

**DOI:** 10.1101/040667

**Authors:** David J. Arenillas, Alistair R.R. Forrest, Hideya Kawaji, Timo Lassman, The FANTOM Consortium, Wyeth W. Wasserman, Anthony Mathelier

## Abstract

**Summary:** With the emergence of large-scale Cap Analysis of Gene Expression (CAGE) data sets from individual labs and the FANTOM consortium, one can now analyze the cis-regulatory regions associated with gene transcription at an unprecedented level of refinement. By coupling transcription factor binding site (TFBS) enrichment analysis with CAGE-derived genomic regions, CAGEd-oPOSSUM can identify TFs that act as key regulators of genes involved in specific mammalian cell and tissue types. The webtool allows for the analysis of CAGE-derived transcription start sites (TSSs) either provided by the user or selected from ~1,300 mammalian samples from the FANTOM5 project with pre-computed TFBS predicted with JASPAR TF binding profiles. The tool helps power insights into the regulation of genes through the study of the specific usage of TSSs within specific cell types and/or under specific conditions.

**Availability and implementation:** The CAGEd-oPOSUM web tool is implemented in Perl, MySQL, and Apache and is available at http://cagedop.cmmt.ubc.ca/CAGEd_oPOSSUM.

**Supporting Information:** Supplementary Text, Figures, and Data are available online at bioRxiv.

## Introduction

The Cap Analysis of Gene Expression (CAGE) technology (Kanamori-Katayama et al. (2011)) has revolutionized our capacity to analyze the promoter and enhancer regions - the c/s-regulatory regions - that are active in cell types and tissues at specific timepoints. Over 1×10^6^ human and 6×10^5^ mouse CAGE peaks were identified through the FANTOM5 project (The FANTOM Consortium and the RIKEN PMI and CLST (dgt) (2014; Lizio et al. 2015)), providing both the precise location of transcription start sites (TSSs) and a quantitative measure of transcriptional activity in the studied samples. It provides the scientific community with an unprecedented opportunity to analyze the *cis*-regulatory regions acting upon the transcription of RNAs from the identified TSSs.

TFs bind to the DNA in a sequence-specific manner at TF binding sites (TFBSs) to regulate gene transcription. Classically, TFBSs are modeled using TF binding profiles, which capture the prefered DNA motifs bound by the TF (Stormo (2013)). Tools such as oPOSSUM3 (Kwon et al. (2012)) and HOMER (Heinz et al. (2010)) provide the ability to perform TF binding profile over-representation analyses on sets of genomic regions to infer the likely TFs acting upon them (for instance, gene promoter regions). Specifically, the tools look for the enrichment of motifs in a set of foreground sequences compared to a set of background ones.

By combining sample-specific CAGE-derived TSSs with TF binding profile enrichment analysis, it now becomes possible to predict the TFs that are likely involved in the regulation of active genes associated with TSSs. We hereby introduce the CAGEd‐ oPOSSUM tool to perform TF binding profile enrichment analysis from genomic regions derived from CAGE data.

This work is part of the FANTOM5 project. Data downloads, genomic tools, and co‐published manuscripts are summarized here http://fantom.gsc.riken.jp/5/.

## Usage and implementation

Through a web interface, CAGEd-oPOSSUM provides functionalities to determine the enrichment of TF binding profiles in a set of user selected CAGE-derived genomic regions by comparing the frequency of predicted TFBSs in the regions versus their frequency in background regions.

**Fig. 1.**
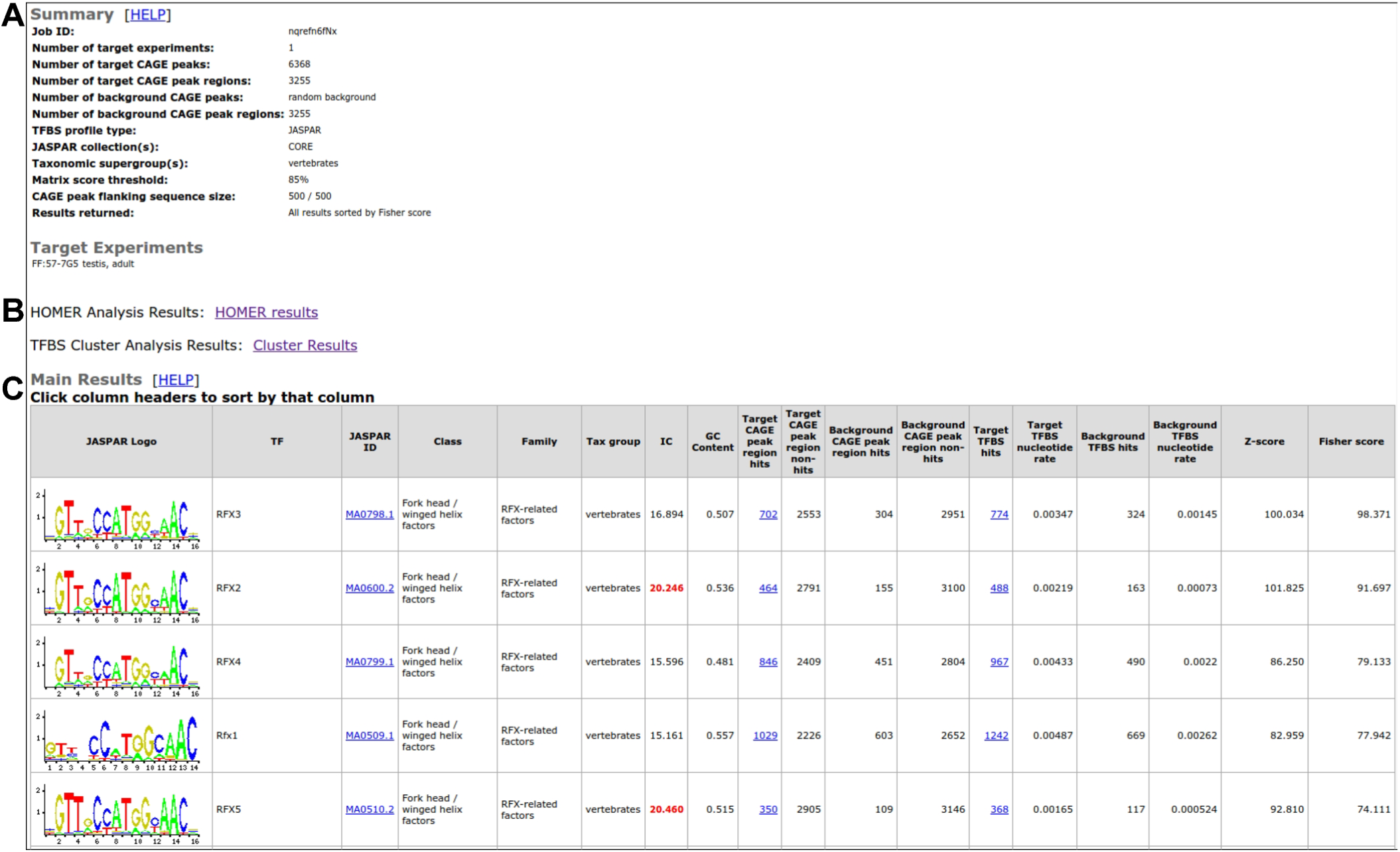
CAGEd-oPOSSUM output example. Screen shot of the CAGEd-oPOSSUM output obtained with the mouse testis, adult (FF:57-7G5) FANTOM5 sample. (**A**) The top of the output page provides a summary of the parameters used by CAGEd-oPOSSUM followed by the list of FANTOM5 samples selected in the foreground. (**B**) It is followed by links to the results obtained using HOMER and the TF binding profile clusters. ( **C**) The bottom of the page provides a table listing the TF binding profile enrichment information (only the top 5 enriched profiles are shown here).

## CAGE-derived genomic regions selection

The user can either provide a set of CAGE peak regions as a BED-formatted file, supply a set of FANTOM5 CAGE peak identifiers (The FANTOM Consortium and the RIKEN PMI and CLST (dgt) (2014)), or select FANTOM5 samples from which CAGE peaks will be selected according to expression level characteristics. To facilitate ease of selection in the latter case, an expandable tree of FANTOM5 samples, based on the corresponding FANTOM5 sample ontology (The FANTOM Consortium and the RIKEN PMI and CLST (dgt) (2014)) derived from cell types (M. Ashburner et al. (2000)), anatomical (Mungall et al. (2012)), and disease ontologies (Smith et al. (2007)), is displayed. The user then filters CAGE peaks based on expression characteristics, proximity to known genes, or overlap with user-provided genomic regions to be used for defining the foreground set of genomic regions around TSSs (see Supplementary Text for details).

A similar procedure is used to select a background set of CAGE peaks. As an additional option for the background, the user may choose to use a random set of regions that is automatically generated to match the %GC composition and length characteristics of the foreground CAGE-derived genomic regions using the HOMER software (Heinz et al. (2010)).

## TFBS prediction and enrichment analysis

Once the set of CAGE peaks is determined, CAGEd-oPOSSUM will extract genomic regions around the corresponding TSSs (for instance using 500 bp up‐ and downstream). Note that overlapping regions are merged before the prediction of TFBSs using BEDTools (Quinlan and Hall (2010)) to avoid counting the same TFBS multiple times. A set of TF binding profiles is then selected from the JASPAR database (Mathelier et al. (2015)) or provided by the user to predict TFBSs in the extracted genomic regions. By default, two statistical tests are employed to determine significance as previously described in oPOSSUM3 (Kwon et al. (2012)): (1) a Z-score that measures the change in the relative number of TFBSs between the foreground and the background sets, and (2) a Fisher score that assesses the number of regions with a TFBS in the foreground set versus the background set. CAGEd-oPOSSUM outputs a result table consisting of a ranked list of TF binding profiles based on the enrichment scores (Fig. [fig:output]). As stated above, the enrichment analysis is performed using oPOSSUM3 statistical tests by default. CAGEd-oPOSSUM additionally allows the user to perform motif enrichment analysis through the HOMER stand-alone package (Heinz et al. (2010)) as a complementary search (Fig. [fig:output]B). Finally, when JASPAR TF binding profiles were used to predict TFBSs, CAGEd-oPOSSUM also performs the enrichment analysis based on TF binding profile clusters derived from profile similarity and defined by the *matrix-clustering* tool in RSAT (Medina-Rivera et al. (2015); Castro‐Mondragon *et al*., in preparation) (Fig. [fig:output]B and Supplementary Text). Specifically, the TF binding profile clusters enrichment analysis is performed following the TFBS Cluster Analysis implemented in oPOSSUM3 (Kwon et al. (2012)).

## Precomputation of TFBSs for time efficiency

As the prediction of TFBSs can be time-consuming, a precomputation was performed from all genomic regions derived from FANTOM5 CAGE peaks using TFBS profiles from the JASPAR database (Mathelier et al. (2015)). All FANTOM5 CAGE peak positions, samples, and expression level data along with the computed genomic regions and predicted TFBSs (see Supplementary Text for details) were stored in a MySQL database. This precomputation allows for the fast analyses of FANTOM5 samples using JASPAR TF binding profiles as the predicted TFBSs are fetched directly from the database to compute the statistical enrichment scores. If the user provides CAGE peak coordinates or TF binding profiles, a similar process is executed in real time.

## Examples of application

As case examples, we tested CAGEd-oPOSSUM on three input FANTOM5 data sets from human primary cell and tissue samples: liver, adult pool1 (FANTOM5 identifier FF:10018-101C9); CD19-positive B-cells, donor 1, 2, and 3 (FF:11544-120B5, FF:11624-122B4, and FF:11705-123B4); and testis, adult, pool 1 and 2 (FF:10026- 101D8 and FF:10096-102C6). Default analysis parameters were used to perform TF binding profile enrichment analyses with both oPOSSUM3 and HOMER for each of these data sets (see Supplementary Text for details).

The most enriched profile predicted by both oPOSSUM3 (using the Fisher scores accounting for the number of genomic regions containing at least one predicted TFBS) and HOMER were associated with TFs known to be involved in the regulation of several biological functions in the corresponding cells and tissues. Namely, HNF4A, ETS factors and RELA, and RFX factors were predicted from the liver, B-cells, and testis samples, respectively (see Supplementary Text, Supplementary Figures, and Supplementary Data for details).

CAGEd-oPOSSUM has been applied to the 1,125 FANTOM5 samples (884 from human and 362 from mouse) and their 418 direct ancestors (241 from human and 56 from mouse) in the ontology trees using default parameters. The results can be found on the CAGEd-oPOSSUM website (http://cagedop.cmmt.ubc.ca/CAGEd_oPOSSUM/results/precomputed/) and links to the specific results are also provided in the corresponding FANTOM5 SSTAR webpages (Lizio et al. (2015)).

## Conclusion

Building upon the large-scale availability of CAGE experiments and established tools (oPOSSUM3 and HOMER) for motif enrichment analysis, we introduced a new computational tool, CAGEd-oPOSSUM, to predict TFs regulating transcriptional events in specific cell types and tissues. CAGEd-oPOSSUM will empower the community with the capacity to highlight specific TFs that may act as key transcriptional regulators in their biological samples of interest when CAGE experiments are available.

